# Aβ42 and Aβ40 oligomers form transient and persistent pores with different time evolutions and toxicities

**DOI:** 10.1101/2022.04.29.490101

**Authors:** Syed Islamuddin Shah, Ian Parker, Ghanim Ullah, Angelo Demuro

## Abstract

In Alzheimer’s disease (AD), formation of harmful self-gating pores formed by the insertion of amyloid beta oligomers (AβOs) into the plasma membrane have been shown to cause disruption of Ca^2+^ homeostasis, leading to neuronal malfunctioning and degeneration. Among different isoforms, the most studied Aβ40 and Aβ42 are also believed to be the most toxic ones. Using single channel imaging, we show that both isoforms can form functionally distinct populations of Ca^2+^ permeable pores, we named transient and persistent pores. The transient pores could be seen only for a few tens of milliseconds, while persistent pores can be observed for more than an hour. However, while the Ca^2+^-toxicity of pores formed by Aβ42Os tend to increase over time by displaying higher open probability and larger Ca^2+^ permeability, pores formed by Aβ40Os show opposite time dependent behavior. We conclude that although both isoforms can form Ca^2+^ permeable pores in the cell’s plasma membrane, pores due to Aβ42Os display worsening Ca^2+^ toxicity over time.

## Introduction

Alzheimer’s disease (AD) is the most prevalent cause of senile dementia, caused by the neurotoxic soluble aggregates of the amyloid beta (Aβ) peptides resulting from abnormal proteolytic processing of amyloid precursor protein (APP) [1-4]. Under suitable condition, soluble Aβ undergo conformational changes that trigger aggregation process, forming oligomeric complexes. Upon interaction with cell membranes, these complexes form cation-permeable pores with functional properties consistent with those of other plasma membrane ion channels [5-11]. By promoting uncontrolled flux across plasma membrane, these pores destabilize the cell’s ionic homeostasis, particularly Ca^2+^, and make the bases for the formulation of the Ca^2+^ hypothesis of AD [5, 9, 12]. Significant progress has been made in establishing the properties of membrane bound Aβ pores [5-11, 13], the molecular mechanism by which these pores contribute to AD [14, 15], and a reduction in their conductivity by various blockers particularly zinc [16-20].

However, a detailed description of how these pores and their toxicity evolve over time, how their gating and conductance vary as the potential difference across cell’s membrane changes, how individual pores respond to antagonists like zinc as they evolve, and many other related questions remain unanswered. In this paper, we address some of these questions by recording the activity of thousands of plasma membrane pores formed by Aβ40Os and Aβ42Os over extended duration with millisecond temporal resolution and sub micrometer spatial resolution. in the absence and presence of zinc, and varying membrane potential.

To study the function of Ca^2+^-permeable ion channels in the cell membrane, we have pioneered an imaging technique called ‘optical patch-clamp’, a massively parallel two-dimensional optical approach, capable of simultaneously and independently monitoring the activity of several thousand channels in their native environment at the single channel and millisecond resolutions [21]. Specifically, we use highly sensitive fluorescent Ca^2+^ indicator dyes in conjunction with total internal reflection fluorescence microscopy (TIRFM) techniques to monitor Ca^2+^ flux through individual Ca^2+^-permeable channels. Binding of Ca^2+^ influx through the channel to cytosolic fluorescent Ca^2+^ indicators generates fluorescent signals (Single Channel Calcium Fluorescent Transients; SCCaFTs) that provide information about channel gating analogous to patch-clamp recording [22-26]. Here we apply this technique to provide the most comprehensive analysis of the formation, gating, and evolution of plasma membrane pores formed by Aβ40Os and Aβ42Os over extended duration.

Our analysis shows that both isoforms can form Ca^2+^-permeable pores, each with two different populations: 1) “<u>transient pores</u>” that can be observed only for a few hundred milliseconds or shorter during multiple recordings, and 2) “<u>persistent pores</u>” that can be observed immobile in the PM for more than hour. However, we observed a clear tendency of Aβ42Os pores to grow over time in number, P_O_, mean open time, and amplitude of Ca^2+^ flux. Conversely, pores formed by Aβ40Os display a tendency to decrease over time in number, mean open time, amplitude. We also present a detailed analysis of the effect of zinc on both pores’ isoforms and their function at different membrane potentials.

## Materials and Methods

Aβs pores’ activity was imaged in the plasma membrane of *Xenopous laevis* oocytes at the single channel level using TIRFM and the data were saved in multi-frame MetaMorph stack files. Stack files were then processed and analyzed using our in-house software called CellSpecks for automatic detection of pores location and extracting gating properties [27]. Further statistical analysis was done in Matlab. Full details of our experimental approach are reported in [9] and are summarized below.

### Preparation and characterization of soluble Aβs oligomers

Solution of 0.5 mg of human recombinant Aβ40 and Aβ42 peptide in 20 μl of freshly prepared DMSO were separately diluted with 480 μl of double-distilled water in a siliconized Eppendorf tube. After 10-min sonication, samples were incubated at room temperature for 10 min and then centrifuged for 15 min at 14,000 g. The supernatant fraction was transferred to a new siliconized tube and stirred at 500 rpm using a Teflon coated microstir bar for 8–48 h at room temperature. Aliquots were taken at intervals and were assayed by pipette application to voltage-clamped oocytes at a final bath concentration of 1 μg/ml. Membrane currents were recorded 20 to 25 min after application in response to hyperpolarization from 0 to −100 mV. Aβ42 preparations incubated for <6 h evoked little or no current, indicating that the monomeric peptide was ineffective at inducing membrane Ca^2+^ permeability. Preparations that evoked currents ≥ 1 μA at membrane potential of −80 mV were used immediately for TIRFM or were stored at − 20°C before use [13].

### Oocyte preparation and electrophysiology

Experiments were performed on defolliculated stage VI oocytes. To visualize Ca^2+^ fluxes, oocytes were injected with Ca^2+^ sensitive dye fluo-4 dextran 1 h before imaging (molecular mass of 10 kD and Ca^2+^ affinity of 3 μM) to a final intracellular concentration of 40 μM. Before TIRF experiments, oocytes were placed in a hypertonic solution (200 mM K aspartate, 20 mM KCl, 1 mM MgCl2, 10 mM EGTA, and 10 mM Hepes, pH 7.2) at 4°C to shrink them so that the vitelline envelope could be manually torn apart and removed using fine forceps. Oocytes were then placed animal hemisphere down in a chamber whose bottom is formed by a fresh ethanol washed microscope cover glass (type-545-M; Thermo Fisher Scientific) and were bathed in Ringer’s solution (110 mM NaCl, 1.8 mM CaCl_2_, 2 mM KCl, and 5 mM Hepes, pH 7.2) at room temperature (23°C) continually exchanged at a rate of 0.5 ml/min by a gravity-fed superfusion system. The membrane potential was clamped at a holding potential of 0 mV using a two-electrode voltage clamp (Gene Clamp 500; Molecular Devices) and was stepped to more negative potentials (−100 mV unless otherwise indicated) when imaging Ca^2+^ flux through Aβs pores to increase the driving force for Ca^2+^ entry in to the cytosol. Solutions containing Aβs oligomers were delivered from a glass pipette with a tip diameter of 25-30 μm positioned near the membrane footprint of the oocyte membrane on the cover glass.

### TIRF microscopy and image acquisition

Imaging was accomplished by using a custom-built TIRFM system based around a microscope (IX71; Olympus) equipped with a 60× TIRF microscopy objective (1.45 NA; Olympus) [28]. Fluorescence excited by a 488-nm laser was imaged using an electron-multiplied charge-coupled device camera (Cascade 128+; Roper Scientific) at full resolution (up to 128 × 128 pixel; 1 pixel = 0.33 μm at the specimen) at a rate of 500 s^-1^. Image data were acquired using the MetaMorph software package (Universal Imaging) and were black-level corrected by subtracting the camera offset. The maximum observed fluorescence signals were small (maximum (F-Fo)/Fo < 2.0, where F is the fluorescence intensity at any time t and Fo is the baseline fluorescence intensity without any channel activity) (see for example Figure 1A1) in comparison to the full dynamic range of fluo-4 ((F-Fo)/Fo >30 in saturating Ca^2+^) and are thus expected to be linearly proportional to Ca^2+^ flux.

**Figure 1.**
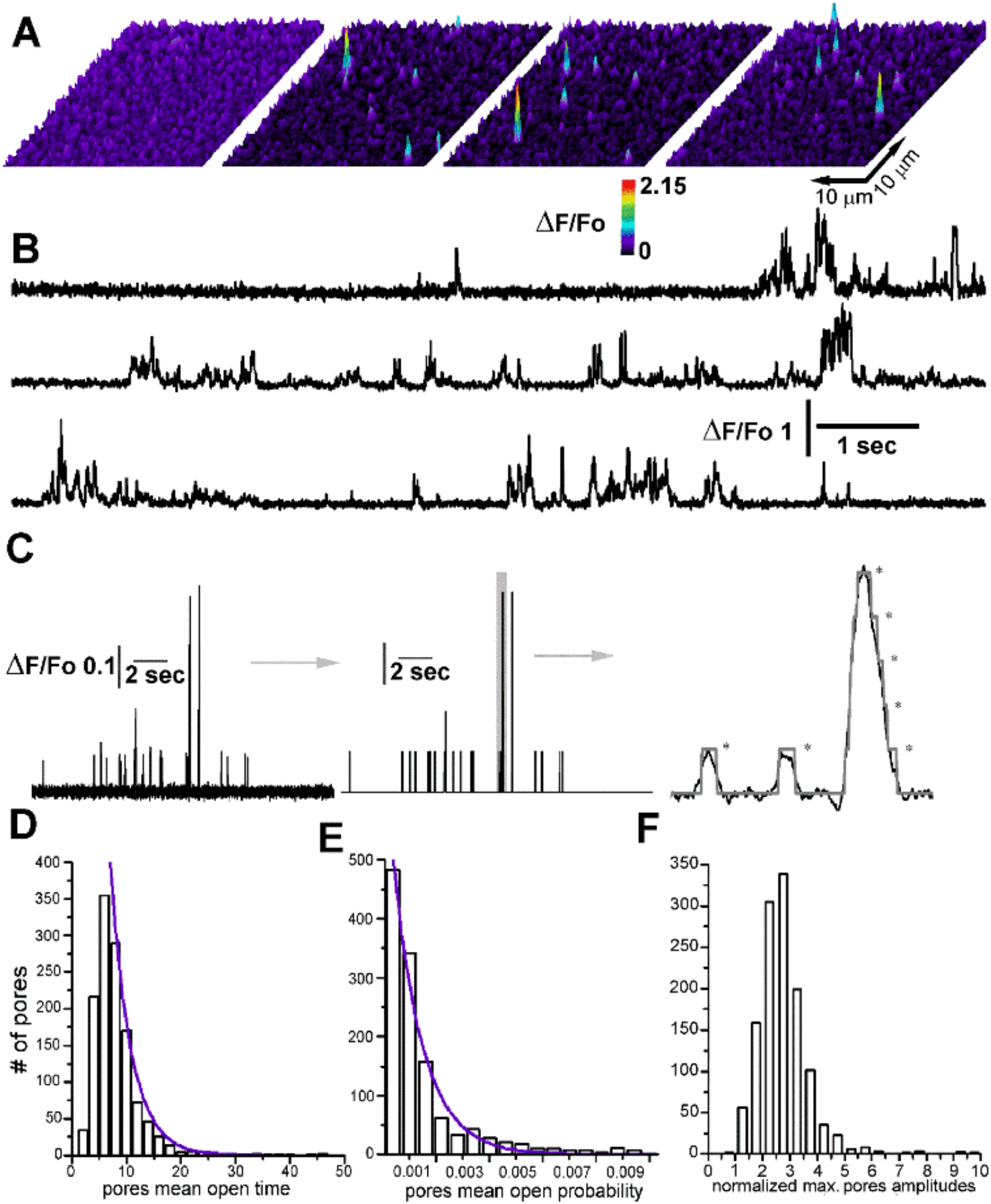
Simultaneous and independent recording of hundreds of functional Aβ42 pores with ms temporal resolution. Bath application of Aβ42 oligomers (final concentration 1 mg/ml) triggers the formation of Ca^2+^ permeable pores in the plasma membrane of *Xenopus* oocytes. **A**) Series of selected single shot images (2 ms exposure time) obtained by TIRFM depicting fluorescent Ca^2+^ transients generated by the opening of Aβ42 pores within a 40×40 μm^2^ of membrane patch of a voltage-clamped oocyte. Frames are captured before (first frame) and during (successive three frames) a voltage step from 0 mV to −80 mV. Increasing [Ca^2+^] is denoted both by “warmer” colors and by height. **B**) Representative fluorescent traces from three different Aβ42 pores showing multiple fluorescence amplitudes corresponding to multiple conductance levels for Ca^2+^ flux through each Aβ42 pore. **C**) A series of steps automatically performed by CellSpecks. Traces are first normalized with respect to resting fluorescence intensity ((F-F_O_)/F_O_) and corresponding idealized traces (center panel) are automatically generated. In right panel, expanded and superimposed fluorescent and idealized traces showing multiple subconductance levels. Idealize traces, are than used to compute distribution of mean open time (**D**), open probability (**E**), and maximum amplitudes for many hundreds of pores in each stack (**F**). Notice that the maximum amplitude is shown in terms of the normalized observed fluorescence intensity. The blue curves are single exponential fits, with decay constants of 9.7 ms (**D**) and 0.0011 s (**E**) respectively. Data are results from 1280 pores from a single stack record.

### Image Processing

As stacks data on the activity of the Ca^2+^ conducting pores contain several thousands of these pores with tens of thousands of events of durations ranging from few milliseconds to many seconds with varying conducting levels, the use of an in-house software package called CellSpecks to process these movie records is essential [29]. CellSpecks is capable of identifying the raw traces of these individual pores, their locations, open and close events along with their durations, mean open (τ_o_) and close (τ_c_) durations, P_o_, maximum flux (amplitude) during each opening event, and the highest Ca^2+^ flux (maximum amplitude) ever observed for a pore during a single stack. Details of how CellSpecks perform these operations and the algorithms it uses to process these movie stacks are described elsewhere [29].

### Pore tracking in space and time

In order to track Aβ pores’ location and activity over extended period of time consecutive image stacks were recorded at the same membrane patch. After collecting several movie stacks (where each stack is up to 30 sec long) at different times after Aβs injection, location maps (x and y coordinates of pixels hosting the pores) of all pores in each movie stack were generated using CellSpecks. To link each pore to a location in multiple stacks, we developed a Matlab routing capable of linking each pore in any stack (using x and y coordinates) with the location in any other stack recorded before or after any given stack. This is achieved by matching the (x, y) coordinates of a given pore in one stack against the coordinates of all pores in other stacks. For example, in a 7 stack experiment, if a pore in stack 3 has coordinates (m, n), we look through stacks 1, 2, and 4-7 to see how many of these other stacks have a pore with the same coordinates. Using this approach, we identified pores which appear only in one stack (named transient pores) and pores that appear in more than one stacks at the same coordinates (named persistent pores). Since CellSpecks also saves time traces and statistics like τ_o_, τ_c_, P_o_, amplitudes of individual opening events, and maximum amplitude (peak amplitude in a time trace) for each pore along with their coordinates, we later analyzed how the properties of transient and persistent pores change over time from one stack to another.

## Results

Formation of self-gating Ca^2+^-permeable Aβs pores in the plasma membrane of specific neurons of AD-affected brains is considered one of the leading mechanisms responsible for the disruption of cellular Ca^2+^ homeostasis[30]. Over time, the uncontrolled Ca^2+^ rise triggers cellular malfunctioning, affecting synaptic function and causing cell death. Using fluorescent Ca^2+^ imaging, we have shown that Ca^2+^-permeable pores formed by Aβ42 oligomers, evoke uncontrolled Ca^2+^ fluxes [9] and suggested that continued evolution of these pores over time may result in the formation of more toxic pores. Here, we provide a high throughput investigation on how the functional properties of these individual pores change over time and compare the functional properties of the two most toxic Aβ isoforms, Aβ40 and Aβ42 in their aggregated forms.

### Imaging Aβ42 pores’ function with high spatial and temporal resolution

Over several years we have developed an imaging technique (optical patch clamping) that permits the gating of hundreds of Ca^2+^-permeable channels to be simultaneously and independently monitored in intact cells with sub-micrometer spatial and millisecond temporal resolutions approaching the accuracy of the electrophysiological patch-clamp [23, 24, 26]. The principle is to monitor cytosolic fluorescence signal generated by Ca^2+^ indicator in proximity of the pore mouth, where large and rapid changes in Ca^2+^ concentration ([Ca^2+^]) occur as the channel opens and closes. This is achieved using TIRFM to excite fluorescence within the ∼100 nm evanescent field. The optical section thus obtained is much thinner (attoliter) than achieved by confocal microscopy, and the planar excitation enables use of highly sensitive, fast (500 fps) ccd cameras. The recordings consist of gray scale movies (often several hundreds of megabytes) generated by TIRFM containing the activity of thousands of Aβs pores over the duration of 20-30 seconds. In each experiment, we recorded several movie files imaging fluorescent activity from a membrane patch (∼ 40×40 μm^2^) of a voltage clamped Xenopus oocyte. As our interest was to correlate changes in Aβs pores’ functioning over time, the stacks are recorded at regular intervals (∼5 mins) staring 20 or 25 minutes after intracellular injection Aβs oligomers. Analysis of this data allows simultaneous and independent investigation of the spatiotemporal activity of thousands of functional Aβs pores present in the membrane patch. As such, a comprehensive analysis of these data has required the development of computational tools allowing automatic detection and analysis of functional pores over time.

Figure 1A shows a series of single frame pictures depicting the intensity profile representation of Single Channel Ca^2+^ Fluorescents Transients (SCCaFTs) generated by these pores detected in a membrane patch of (40×40 μm^2^) of a voltage clamped oocyte. The color-coding represents amplitudes where warmer color indicates larger increase in cytosolic Ca^2+^ concertation due to the pore, which also provides location of each functional pore at the time of recording. In Figure 1B, we show selected Ca^2+^ fluorescence traces from individual region of interest positioned centered over a pore location normalized with respect to the background fluorescence (ΔF/Fo) extracted from a 30 second video recording using CellSpecks. In Figure 1C a series of panels describing consecutive steps operated by CellSpecks. First, after removing the noise from row data (left trace), an idealized trace is generated indicating the conductance level (closed state, first open state (state 1) with lowest conductance, second open state with conductance double that of state 1, and so on) in which the pore is gating as a function of time (central trace). Right panel is a zoomed-in to highlight the multi-conductance levels (indicated by stars) in which individual pores can gate. Each stack contains many hundreds, sometimes thousands of pores with varying *τ*_o_, P_o_, and amplitudes which are automatically analyzed and plotted by CellSpeck. In Figure 1D-E mean open time, open probability, and amplitudes distribution for pores detected in this specific stack are shown.

In order to investigate the evolution of Aβs pores over extended period of time, we recorded multiple consecutive stacks at the same membrane patch. We then linked each pore location (x and y coordinates in a sub-pixel pores’ location map) generated by CellSpecks using our Matlab routine capable of linking each pore in any stack with the location in any other stack recorded before or after that stack. In Figure 2A a TIRFM image (left panel) depicting a 40×40 μm^2^ membrane patch of a voltage clamped *Xenopus* oocyte exposed to Aβ42 oligomers from which 7 consecutive stacks were recorded. Continuous formation of new pores was detected at random location in the membrane patch. Selected fluorescent traces from four different locations in the membrane patch, corresponding to the location of four different Aβ42 pores location detected during the 7 consecutive stacks. CellSpecks allows automatic allocation of pores coordinate with super resolution capability, providing pore coordinates with sub-μm resolution. Overlapping the maps of pore locations over consecutive stacks revealed that some pores were present in only one stack whereas others could be observed during the full duration of the experiment as shown in Figure 2A where a pore activity was observed for 6 consecutive stacks.

**Figure 2.**
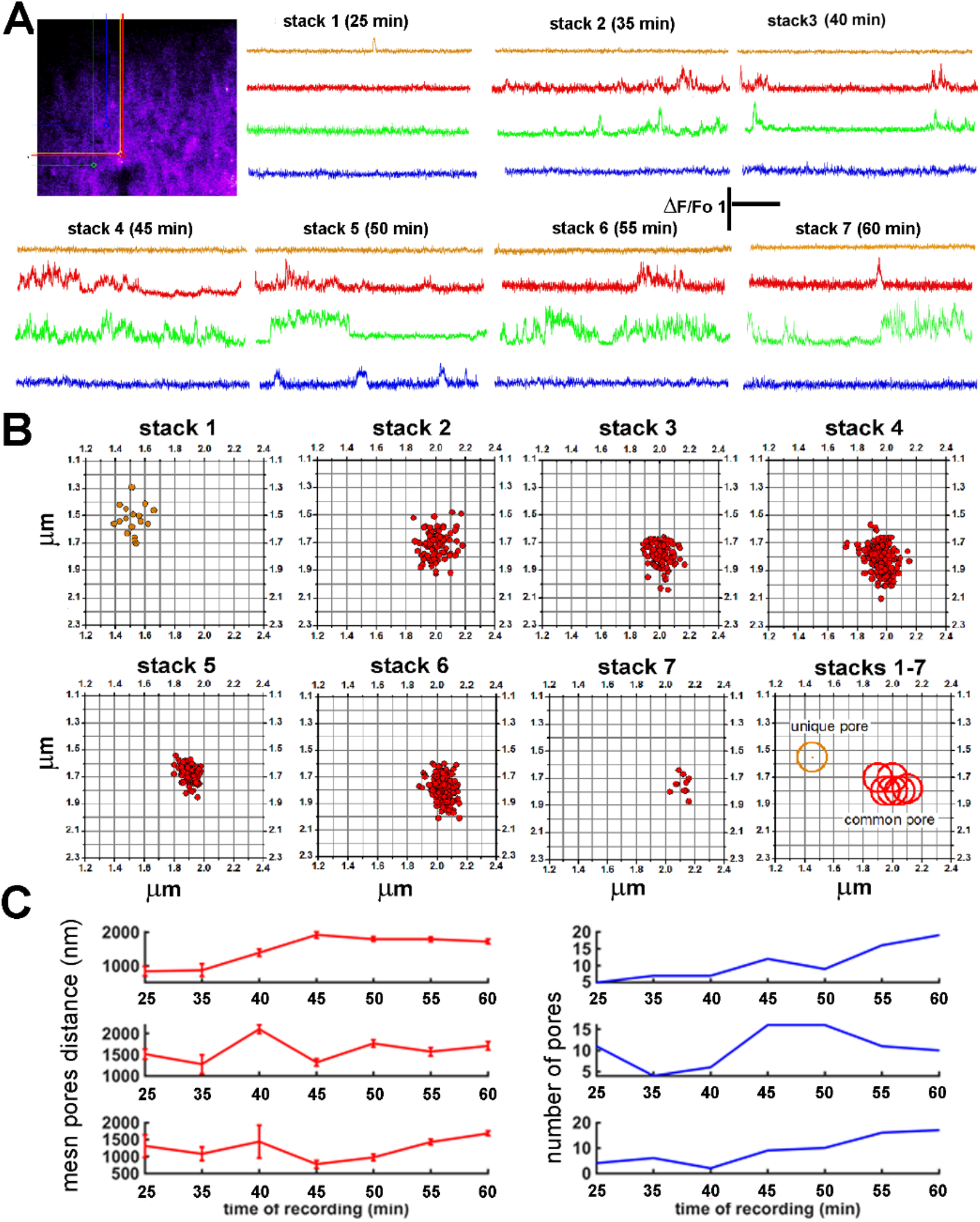
Tracking Aβ42 pore location with sub mm resolution during multiple consecutive recording. **A**) TIRF microscopy image (left panel) depicting a 40×40 μm^2^ membrane patch of a voltage clamped *Xenopus* oocyte exposed to Aβ42 oligomers from which 7 consecutive stacks were recorded. Continuous formation of new pores was detected at random locations in the membrane patch. Selected time-dependent fluorescent traces from four different locations in the membrane patch, corresponding to the location of four different Aβ42 pores as indicated in the maps. Traces displaying different single channel behavior during the recording of seven consecutive stacks, with pores active only in one stack and other observed during the full duration of the experiment, in this case visible for 6 consecutive stacks. **B**) Super resolution tracking of 2 neighboring pores (top two traces panel **A**) in a 1.2 μm x 1,2 μm membrane patch. Top panel, from left to right, location of active pores from stack1 to stack7. Number of dots in each panel represents the location of the single SCCaFTs each time the pore was active. A maximum radius of 100 nm (sub pixel) was assigned for the pore tracking location and only centroids displaced within 200 nm (distance between two centroid) in consecutive frames are assigned to the same pore. The bottom right panel depicts superimpose centroid locations for each pore, i.e. geometric center for all the single SCCaFts during consecutive recordings. **C**) Analysis of 5×5 μm^2^ membrane patches shows that Aβ42 pores do not overlap and their local density changes over time. Panels on the left depict the mean distance of all pores around a common pore revealing low probability of overlap as the mean distance is much larger than the spatial resolution of our imaging system. Right panels show the number of pores indicating that density of pores changes with time. Regions with three different pore densities were selected for this analysis.

To demonstrate that our pipeline can clearly separate nearby pores at sub pixel resolution and distinguish pores appearing in one stack from those appearing in multiple stacks, we performed super resolution spatial tracking using a maximum radius of 100 nm (sub pixel resolution) around the centroid of individual SCCaFTs. Only centroids displaced less than 200 nm in consecutive stacks were assigned to the same pore. This analysis is shown in Figure 2B, where centroids of SCCaFTs generated by the opening of two nearby pores imaged during seven consecutive stacks are shown. Each panel is a 1.2×1.2 μm section of the same area of the image field and each dot represents the centroid (geometric center) of fluorescent signal when the pore was open in a frame. In the first panel, we show the location of the pore that was only active in the first stack. Each orange dot represents the location of the fluorescent centroid when the pore was active in a given frame. Thus, the number of dots represents the number of frames in which this pore was active. The red dots in the second panel have been assigned to a different pore and was detected in the next consecutive six stacks. Next, we calculated the geometric center of all the centroids in a given stack, which represents the location of the pore in that stack. In the right bottom panel, the location of the pores in each stack are reported with a circle having a radius of 100 nm. The pore appearing in one stack (one orange circle) can be clearly distinguished from the pore that persists in six stacks (six overlapping red circles). The overlapping of circles also shows that the second pore stays at the same location (within 100 nm) throughout the experiment. Corresponding time dependent fluorescent traces recorded from these two locations are shown by orange and red traces in Figure 2A, respectively.

To rule out the overlap of multiple pores, we analyzed pore density in three randomly selected 5×5 μm^2^ membrane patches, each around a pore that persisted throughout the experiment (seven stacks). We calculated the mean distance of all pores around a common pore in the 5×5 μm^2^ membrane patch. The mean distance between pores in each patch mostly remained larger than 1 μm, much larger than the spatial resolution of our imaging system, suggesting that the probability of different Aβ42 pores in our experiments to overlap is low (Figure 2C, left panels). Right panels show the corresponding number of pores within these three 5×5 μm^2^ patches, demonstrating that while the number of pores continue to change (mostly increasing), the mean distance between the pores remains well above the spatial resolution of our system.

### Tracking single pore functions revealed the formation of transient and persisten pores with increasing Ca^2+^ toxicity over time

In Figure 3, we show the cumulative results obtained by analyzing a series of 5 experiments including multiple imaging stacks acquired from 5 different oocytes exposed to 1μg/ml of Aβ42 oligomers. Experiments were performed as indicated in Figure 2, acquiring ∼30 sec video sequence every five minutes for up to 8-16 stacks. In the analysis reported in Figure 3 some stacks were not included, due to small changes in focus, producing blurry images that prevent automatic analyses by CellSpecks. Additionally, the analysis of these experiments was limited to 8-13 consecutive stacks recording as increasing bleed through in fluorescence signals between nearby pores, as a result of increased pores density and larger single channel Ca^2+^ fluxes prevent automated analysis. In Figure 3A-C are plots showing total number of pores, their averaged number, and percent change calculated for each corresponding time of recording. The high variability in the total number of pores at each recording time (Figure 3A) was due to the exclusion of stack in some of the experiments and variability in the pores number in each experiment. However, when the number of pores was averaged for each recording time (Figure 3B) or expressed as percent increase over the pores number obtained in each first recording stacks (Figure 3C) the growing trend in pore number over time is more evident. Similar trend was observed for the percent change in mean open time (Figure D) and the percent change in open probability over time (Figure 3E). In Figure 3F, we show the averaged maximum amplitudes (maximum Ca^2+^ fluorescence observed in a time trace) of pores measured in each stack for each recording time, displaying a gradual increase in pores’ maximum amplitude over 60 minutes. For each recording the corresponding location map for all the pores were automatically recorded. Overlapping of these pore location maps obtained during each single experiment performed on the same membrane patch revealed that some pores were present in many consecutive stacks (persistent pores) whereas others could be detected only in one stack (transient pores) during the entire experiment consisting of several stacks.

**Figure 3.**
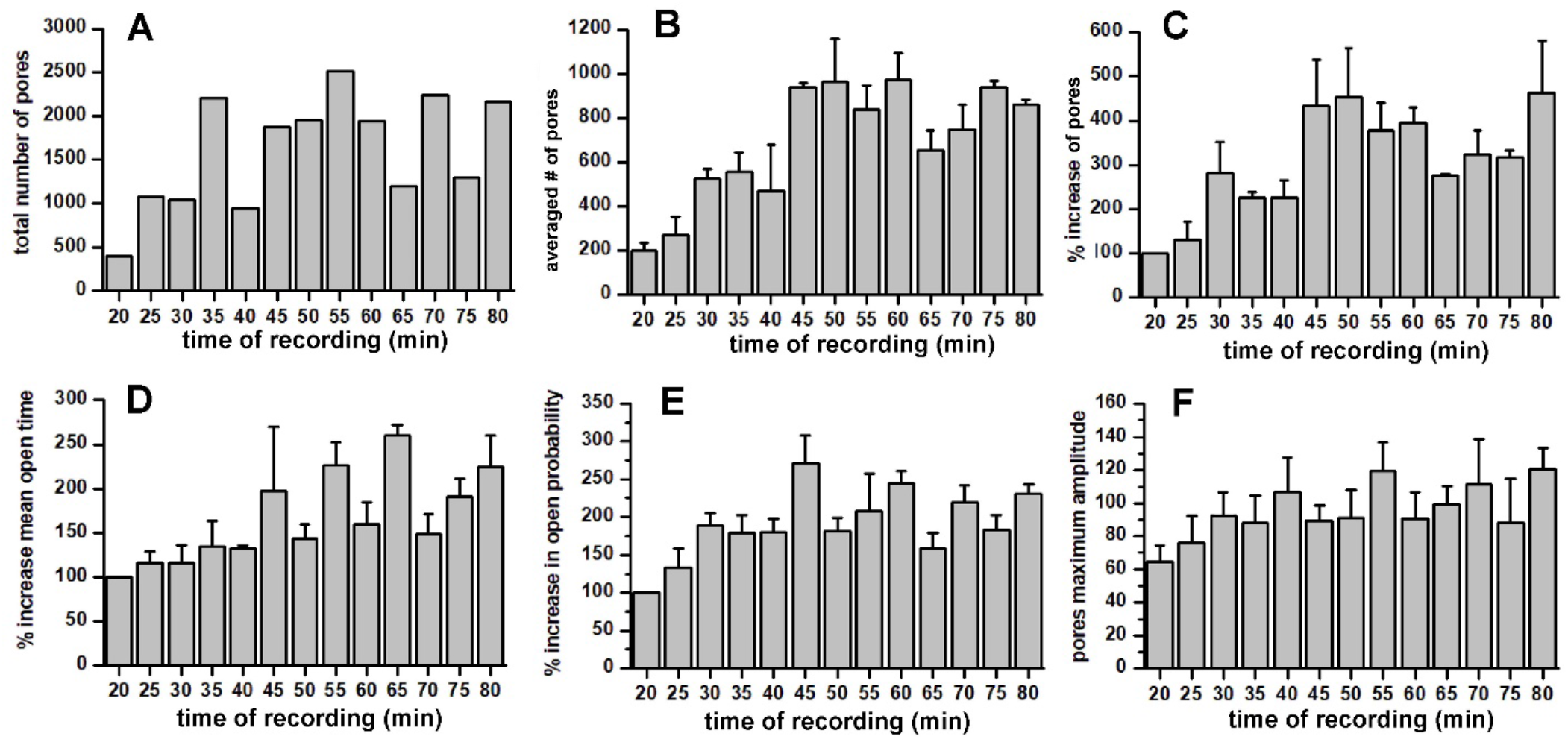
Time dependent changes in Aβ42 pores’ activity. **A**) Cumulative plots displaying the total number of pores detected during the recording of consecutive stacks acquired at the time after Aβ42 exposure as indicated by the horizontal axis. Data are obtained from 5 different oocytes and each column represents the sum of the pores recorded at the indicated time from multiple stacks. **B**) Same data as in **A** plotted as the average number of pores (pores per oocyte) recorded at the corresponding recording time showing a clear increase in the number of pores over time. **C**) Same data replotted as normalized with respect to the first recording, showing a percent increase in pores number over time. **D**) Pores mean open time increases during consecutive recordings. The calculated values of mean open time for each recording during each experiment were normalized to the value obtained from the first recording for each respective experiment. Values for each time recording are then averaged and plotted as a single column. **E**) Corresponding increase in mean open probability normalized to the first recording time showing a time dependent increase in open probability. **F**) Absolute values of maximum pores amplitude (βF/Fo) increase over time. Each column represents the averaged value of maximum pore amplitude calculated for each recording performed at the same time after Aβ42 application. Data in each column depict the mean values +/- SE of measurements from two to four experiment from three frog donors.

To perform in-depth analysis of the behavior of these two different populations of pores, we selected a single experiment that included seven consecutive stacks at the same membrane patch. In Figure 4A, the resulting statistical analysis is reported in a series of plots where the number of pores, corresponding mean open time, mean open probability, mean maximum amplitudes, and mean close time are reported as a function of time for all pores in the membrane. Each column represents the average value ± standard error. In Figure 4B, a series of panels depicting the 40×40 μm^2^ patch of the oocyte membrane show the corresponding locations of the Aβ42 pores analyzed in Figure 4A. Maps depict pore locations during the seven consecutive stacks recorded every 5 minutes. From left to right, the first map (Figure 4B, top left) shows pores locations in the first stack, second panel shows the resulting overlapping maps of pore locations detected in stack 1, stack 2 and stack 3. Third panel is generated by the overlapping of pore maps generated from stack1 to stack 5, and fourth panel shows the overlapping of all maps generated for pore locations in stack 1 to stack 7. Pore locations for each stack are indicated by dots with selected color and shape for each stack. Fifth and sixth panels display two consecutive magnifications (as indicate by the arrows) of the smaller region in panel four (stack1-7). On panel six (map1-7 zoom 2), pores detected only once or multiple times can easily be identified by eye as is clear from the presence of one single symbol or multiple overlapping of color-coded symbols, respectively. Analysis of the pore location within these seven maps, revealed two population of pores. One, we named *transient* pores, which are detected only in one stack throughout the entire experiment. The second population, we named *permanent* pores, are visualized at the same location (within 100 nm resolution) in multiple stacks and can be visualized in up to seven consecutive recording stacks. A summary of this analysis is shown in Figure 4C where the color-coded table (wormer blue representing larger number of pores) illustrates the total number of pores observed in each stack (diagonal elements) and the number of pores that have been observed in multiple stacks (off-diagonal elements) during the experiment. Horizontal and vertical axes represent the time at which each stack have been recorded after application of Aβ42 oligomers. For example, 16 out of the total 244 pores observed in the first stack (25 min after Aβ42 application) are also observed in the second stack recorded after the first stack. Similarly, 89 out of 477 total number of pores observed in the second stack (35 min) are also observed in the third stack recorded after the second stack, and so on. The table on the right shows the number of *transient* pores detected in any single stack and included in the total number of pores in the diagonal elements of the main table on the left.

**Figure 4.**
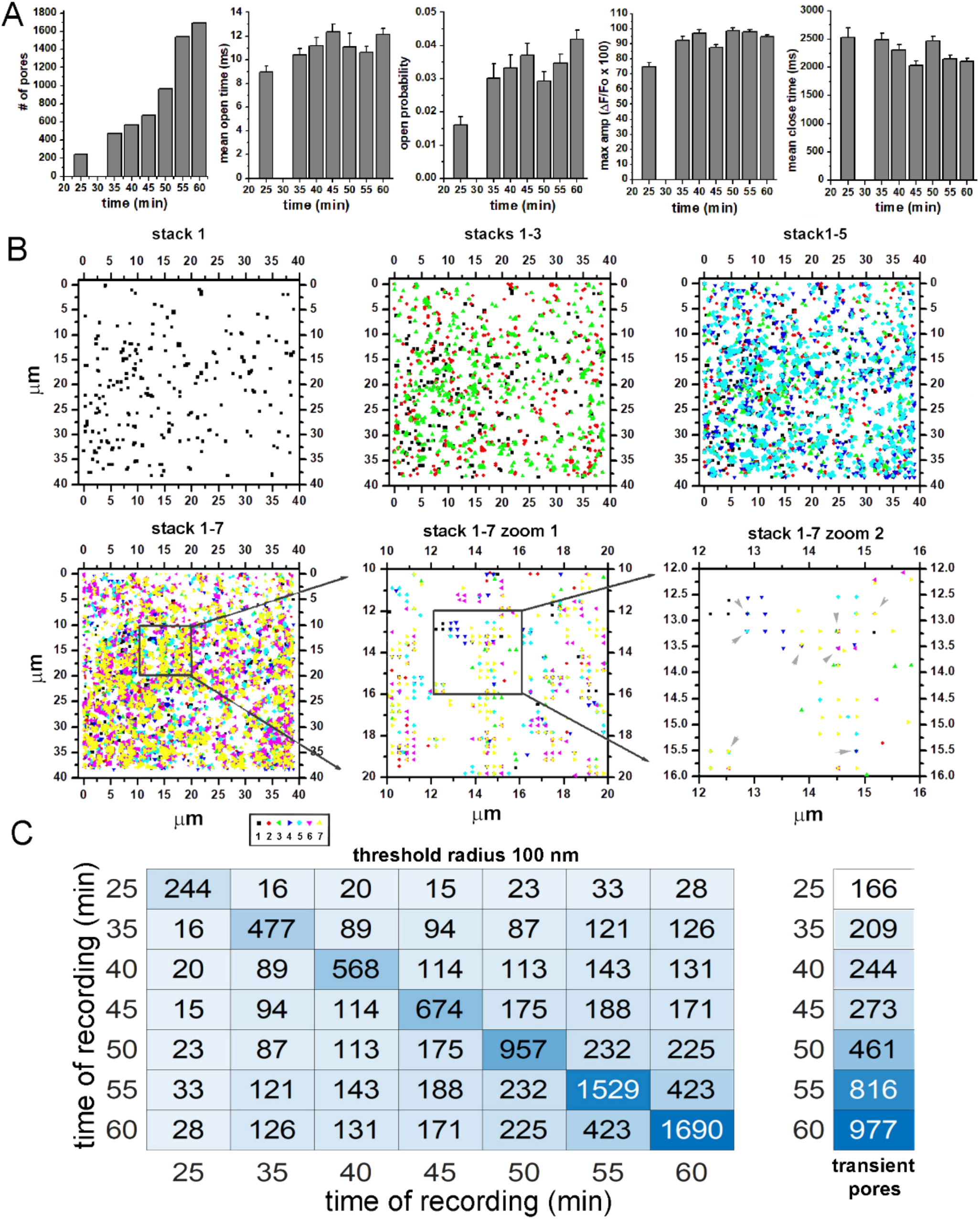
Overlapping of consecutive maps of pore locations reveals two different functional population of Aβ42 pores. A single experiment consisting of 7 consecutive stacks obtained from the same imaged field was analyzed by tracking each pore’s location over time independently and with sub-μm resolution. **A**) The number of pores and the corresponding mean open time, mean open probability, mean maximum amplitudes and mean close time as a function of time. **B**) Overlapped location maps of Aβ42 pores analyzed in **A**, within a 40×40 μm^2^ of membrane patch. The stack numbers to which the maps belong are indicated on top of each panel (also indicated by color and shape of symbols). The last two panels display two consecutive magnifications (as indicate by the arrows) of the smaller region in panel four (stack1-7). **C)** Color coded table (warmer blue representing larger number of pores) illustrating the total number of pores observed in each stack (diagonal elements) and the number of pores that have been observed in multiple stacks (off-diagonal elements) during the experiment. Horizontal and vertical axes represent the time at which each stack was recorded after application of Aβ42 oligomers. For example, 16 out of the total 244 pores observed in the first stack (25 min after Aβ42 application) are also observed in the second stack recorded after the first stack. Similarly, 89 out of 477 total number of pores observed in the second stack (35 min) are also observed in the third stack recorded after the second stack, and so on. The right-most column shows the number of pores that are unique in any single stack. Note that the total number of unique and common pores in a given row can be larger than the total number of pores (diagonal elements in the left table) because some pores were observed in more than two stacks.

Next, we investigated independently the evolution over time of these two populations of pores. The results of this analysis are shown in Figure 5, where values for mean open time for transient pores, permanent pores, and total pores (transient and persistent combined) are plotted separately (Figure5A). Their corresponding maximum pore amplitudes and open probabilities are shown in Figure 5B and 5C, respectively. From this analysis it is evident that *persistent* pores tend to display higher mem open time, maximum amplitudes, and open probability over time when compared to *transient* pores. This difference in pores functioning over time was particularly evident in the open probability.

**Figure 5.**
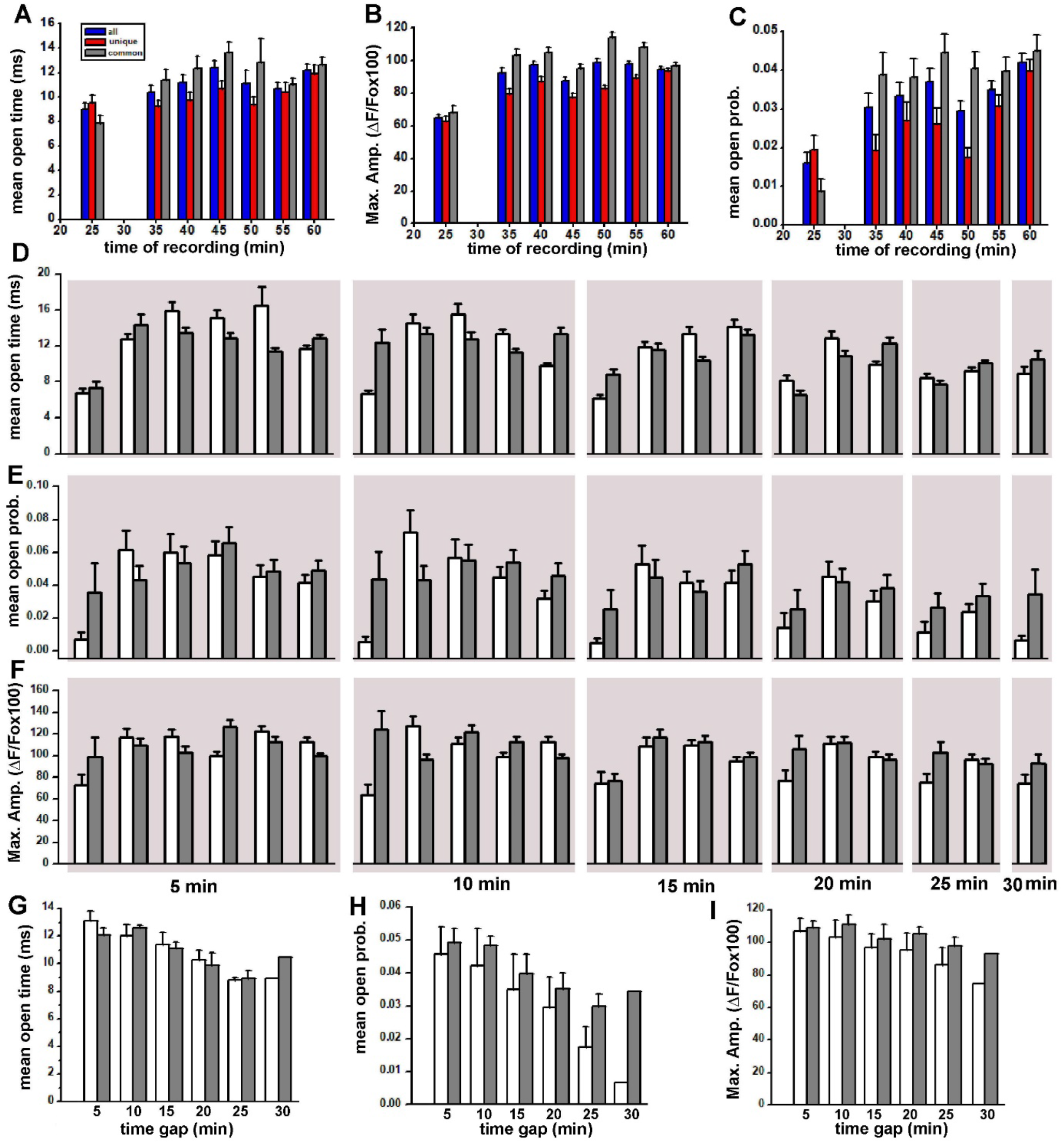
Tracking transient and persistent Aβ42 pores’ function during multiple stacks. The ability to track individual pores with a threshold of radius of 100 nm with high level of certainty allows accurate classification of transient and persistent pores for a separate analysis. **A**) The averaged values of the mean open time for all pores (transient and persistent) (blue bars), transient pores (red bars), and persistent pores (gray bars) observed during each of the 7 consecutive stacks. **B, C**) The corresponding averaged maximum pore amplitude and averaged open probability. **E-F**) Average statistics of all the pores that are common in any two stacks: **D)** mean open time, **E)** open probability, and **F)** max amplitude. White bars represent mean open time, open probability, and max amplitude of the pores that appear in the earlier stack whereas gray bars represent the corresponding measurements for the same pores that appeared in the later stack. First six pairs of bars represent pores that appear in two consecutive stacks separated by 5 minutes (time indicated in **F**) such as stacks recorded at 25 and 30 minutes, 35 and 40 minutes. The next set of bars represent pores that appear in two stacks separated by 10 minutes (e.g. 25 and 35 minutes) and so on. Plots in **G, H** and **I** display the average changes in pores’ parameters detected in any two stacks combination with growing time gaps. For example, all the white bars in top-left panel in **D** are averaged to get the first white bar (5 minutes) in **G**.

To gain a deeper insight into the evolution of persistent pores, we investigated how the gating properties of these pores in one stack compare with their properties in another stack recorded latter in the experiment. To do this, we enumerated stacks in increasing order of time at which they were recorded after exposing the cell to Aβ42 oligomers and generated all possible combinations of stacks. For example, in an experiment with two stacks (say 1 & 2; labeled as 12), we have only one combination to search for pores that appear in both stacks. For an experiment with three stacks (1, 2 & 3), we have three possible combinations (1 & 2, 1 & 3, or 2 & 3, labeled as 12,13, and 23) to look for pores appearing in two stacks, and so on. Next, we sorted the lists of pores appearing in these different combinations of stacks according to the time separating the two stacks in a combination. Finally, we calculated the averaged mean open time, averaged open probability, and averaged maximum amplitude of all pores in the first and second stack in the combination separately, and report our results in Figure 5D-F. For example, in Figure 5D-F, results from combinations with an inter-stack interval of 5 min (combinations 23, 34, 45 etc.) are included in the first set of plots, where the empty bars represent the average statistics from the first stack and the filled bars represent from the following stack. Pores in combination such as 13 and 24, separated by 10 min interval are included in the next set, and so on. The inter-stack time for each column of panels is indicated at the bottom in Figure 5F. Note that in this experiment with seven stacks, there are 21 combinations of two stacks (6 combinations with inter-stack interval of 5 min, 5 combinations with inter-stack interval of 10 min, and so on). We combined the results from all possible combinations in Figure 5G-I. For example, the empty bar at 5 min is the average of all 6 empty bars in Figure 5D left panel. Similarly, the filled bar at 5 min is the average of all 6 filled bars in Figure 5D left panel. Overall, the activity of the persistent pores is higher in the following stack (at later time) than in the first stack. With a few exceptions, this increased activity of permanent pores at later time is a generalized behavioral trend that we observed in all our experiments. We also analyzed persistent pores in combinations of 3 stacks to (e.g. pores observed in stacks 1, 2, and 3) all the way up to the combination of all 7 stacks (pores observed in all 7 stacks), and found a similar trend (not shown).

### The amplitudes of SCCaFTs due to Aβ42 pores increase linearly with increasing membrane depolarization

SCCaFTs due to Aβ42 pores were clearly visualized at negative potential such as −80 mV. The amplitudes of these events gradually decreased at less negative potentials, almost vanishing as membrane potential was shifted to 0 mV. This is expected for Ca^2+^ permeable ion channels when the driving force for Ca^2+^ influx is increased, and confirms the extracellular source of Ca^2+^. However, it is not clear if the negative membrane potential only affects the Ca^2+^ flux i.e., fluorescence amplitudes, or also affects the gating properties of Aβ42 pores. To answer this question, we monitored the activity of Aβ42 pores under different membrane potentials. Figure 6A top row depicts the time-dependent Ca^2+^ fluorescent intensity generated by averaging the fluorescence traces of all pores that appeared persistently at the same locations in the cell membrane at membrane potentials of 0 mV, −40 mV and −80 mV as indicated by the bars above each trace. The next four rows in Figure 6A show traces obtained from four different 0.6 × 0.6 μm^2^ regions of interest, depicting SCCaFTs at the same four pore locations during the recording of the three stacks. These traces clearly show that not only the amplitudes but the durations of SCCaFTs also change as the membrane potential changes. The averaged fluorescence amplitude of SCCaFTs from all Aβ42 pores as a function of membrane potential is sown in Figure 6B. The fitted regression line extrapolates to 0 at about 32 mV, confirming the typical reversal potential for Ca^2+^ in oocytes. Figure 6C and 6D depict the average values of the mean open probability and mean open time of all pores in the membrane patch at each respective membrane potentials.

**Figure 6.**
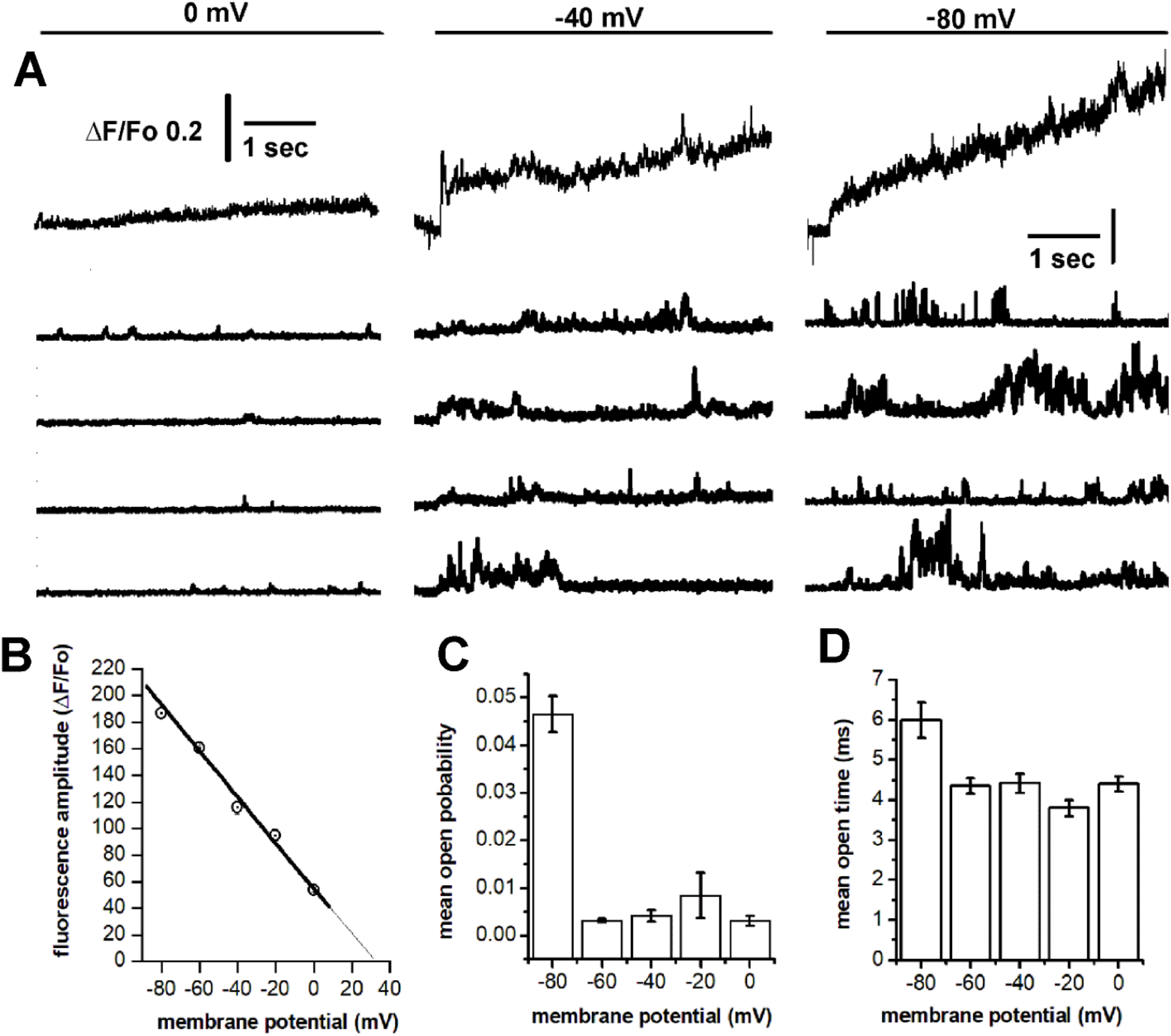
Effect of transmembrane potential on amplitude and gating properties of Aβ42 pores. Recording at different membrane voltage steps are obtained from the same membrane patch. **A**) Traces in top row shows time-dependent Ca^2+^ fluorescent recording obtained by averaging all the fluorescence traces generated by positioning a 2×2 pixels region of interest on each persistent Aβ42 pore location identified in the membrane patch (40×40 μm) at the respective membrane potentials (bars at the top). Four bottom rows show four selected traces depicting SCCaFTs at the same four pore locations recorded at three different membrane voltages. **B**) Mean SCCaFTs amplitude of all pores in each stack as function of membrane potentials. Regression line extrapolate to zero at a potential of about 32 mV. **C**) Plot displaying the respective mean open probability for all pores recorded in each stack and **D**) the corresponding averaged mean open time.

### Zinc inhibits Aβ42 pores reversibly

Zinc ions have been shown to inhibit Aβs pores function, reducing the total Ca^2+^ influx through the cell membrane [14, 16-19]. To quantify the effect of zinc at the single pores level, we performed three independent experiments where the activity of Aβ42 pores were first imaged in the absence of zinc, and after addition of 200 μM at the same membrane patch. The cumulative results of these experiments are shown in Figure 7. A series of representative fluorescent traces obtained from 7 regions of interest (0.6×0.6 μm^2^) positioned at the top of 7 Aβ42 pores’ fluorescence centroids recorded before zinc addition are shown in Figure 7A (left panel). Corresponding fluorescent traces recorded at the same locations in the same membrane patch after addition of 200 μM zinc are shown in the righ panel, where fluorescent transients were strongly inhibited in both, amplitudes, and duration. Figure 7B shows an expanded view (grey boxes in Figure 7A) of fluorescent changes, depicting changes in the single pore function after zinc application. Single channel analysis of the pores recorded before and after zinc application reveals strong inhibitory effect on the number of detectable functional pores after addition of 200 μM zinc (Figure 7C). We also observed considerable reduction of pores maximum amplitudes (Figure 7D) and number of opening events (Figure 7E) after zinc application. However, the average number of events recorded per pore for pores that were still active in the presence of zinc marginally increased (Figure 7F). Overlapped distributions of the measured mean open time before (blue bars) and after zinc application (red bars) display a clear reduction in mean open time caused by zinc (Figure7G). Inset in Figure 7G compares the two distributions after normalization. As shown in Figure 7H, a strong reduction in the Aβ42 pores’ open probability was also observed after zinc addition. Figure 7I shows the corresponding distributions of the mean close times before and after zinc application.

**Figure 7.**
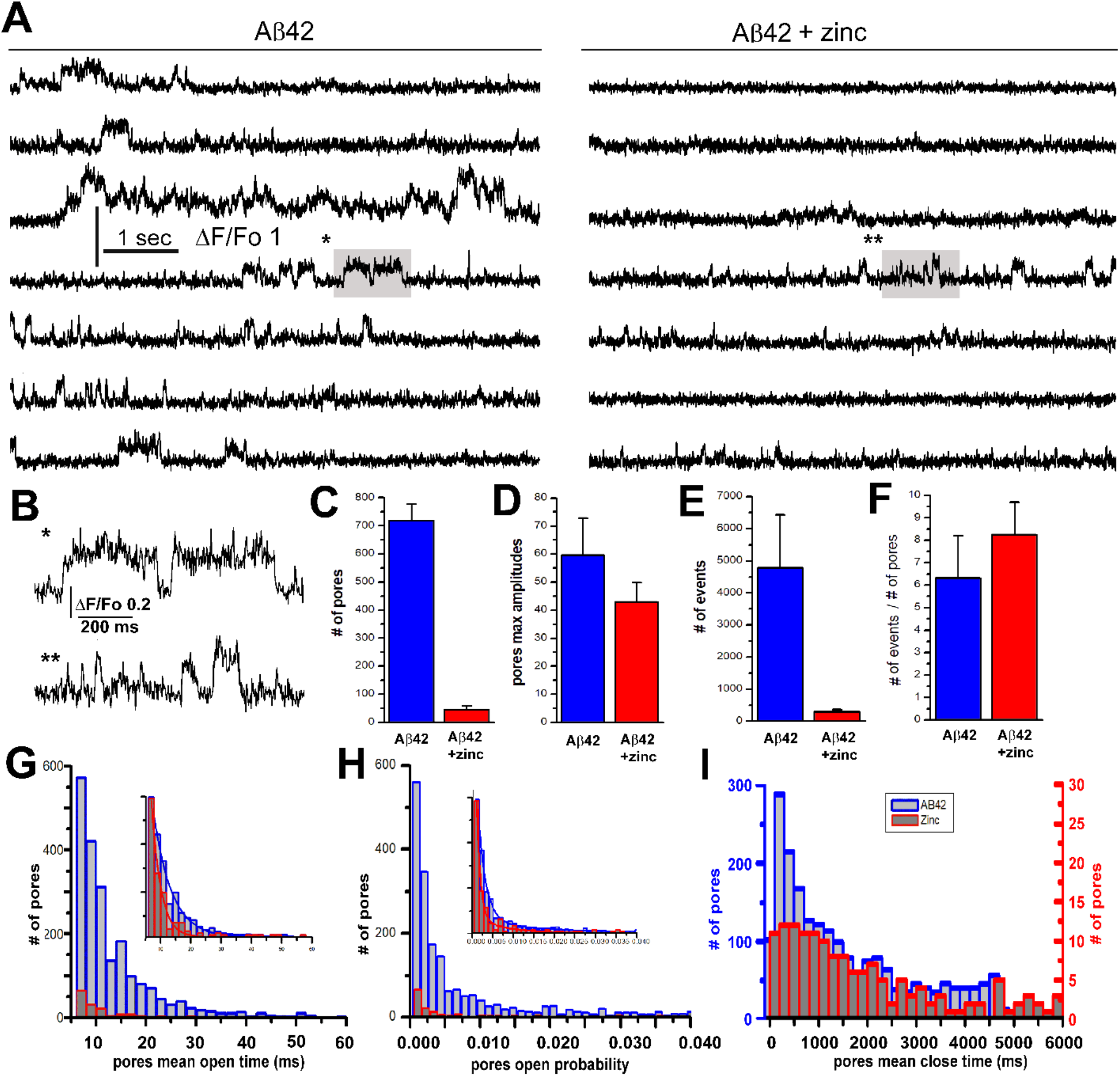
Aβ42 pores’ activity is strongly inhibited by zinc. **A**) Series of representative traces showing fluorescence activity from seven different pores recorded in the absence (left panel) and in the presence of extracellular zinc (right panel) from a single oocyte. Recordings are obtained at about 40 minutes after Aβ42 application. **B**) Expanded sections of fluorescence traces highlighted in the fourth row, showing the reducing effect of zinc on the open time for the same Aβ42 pore. **C**) The mean number of Aβ42 pores from 3 different experiments obtained during application of Aβ42 oligomers alone (blue bars) or when applied together with zinc (red bars). **D** and **E** are the corresponding averaged maximum amplitudes and the average number of events in all three experiments. **F**) Corresponding number of events per pores before and after zinc co-application. **G**) Mean open time distribution for all Aβ42 pores recorded before (total of 2340 pores) (blue bars) and after (red bars) zinc application. Inset shows normalized distribution of the main plot. **H**) Corresponding open probability distribution of all pores before and after zinc application and (inset) normalized distribution. **I**) The corresponding distributions mean closed times of all pores.

To further investigate the modality of zinc inhibition on Aβ42 pores, we imaged Aβ42 pores’ activity before application, in the presence of, and after the removal of zinc in the same patch of cell’s membrane. Location maps of functional Aβ42 pores obtained during a single experiment are shown in Figure 8A, where the map on the left displays pore locations in the presence of Aβ42 oligomers alone. After zinc addition, the resulting number of detected pores are greatly reduced as shown in the center map. After zinc removal, the number of detectable Aβ42 pores increased again reaching even higher numbers as compared to the first recording (map of right). Panels in Figure 8B show the corresponding color-coded (intensity profiles) representation of the temporal evolution of the fluorescence signals due to all pores during the three recordings. Figure 8C shows the total number of pores during each recording conditions, showing strong reduction of detected pores in the presence of zinc. Consistently, reduction in mean open time and open probability was also observed when zinc was present in the bathing solution (Figure 8D, E). These reductions were reverted after removal of zinc, particularly for the mean open probability that increased to higher averaged values as compared to the measurements obtained in the initial recording before the application of zinc. We also observed a moderate reduction in the average mean close time (Figure 6F) and a reduction of ∼ 30% in the averaged maximum amplitude in the presence of (Figure 6G).

**Figure 8.**
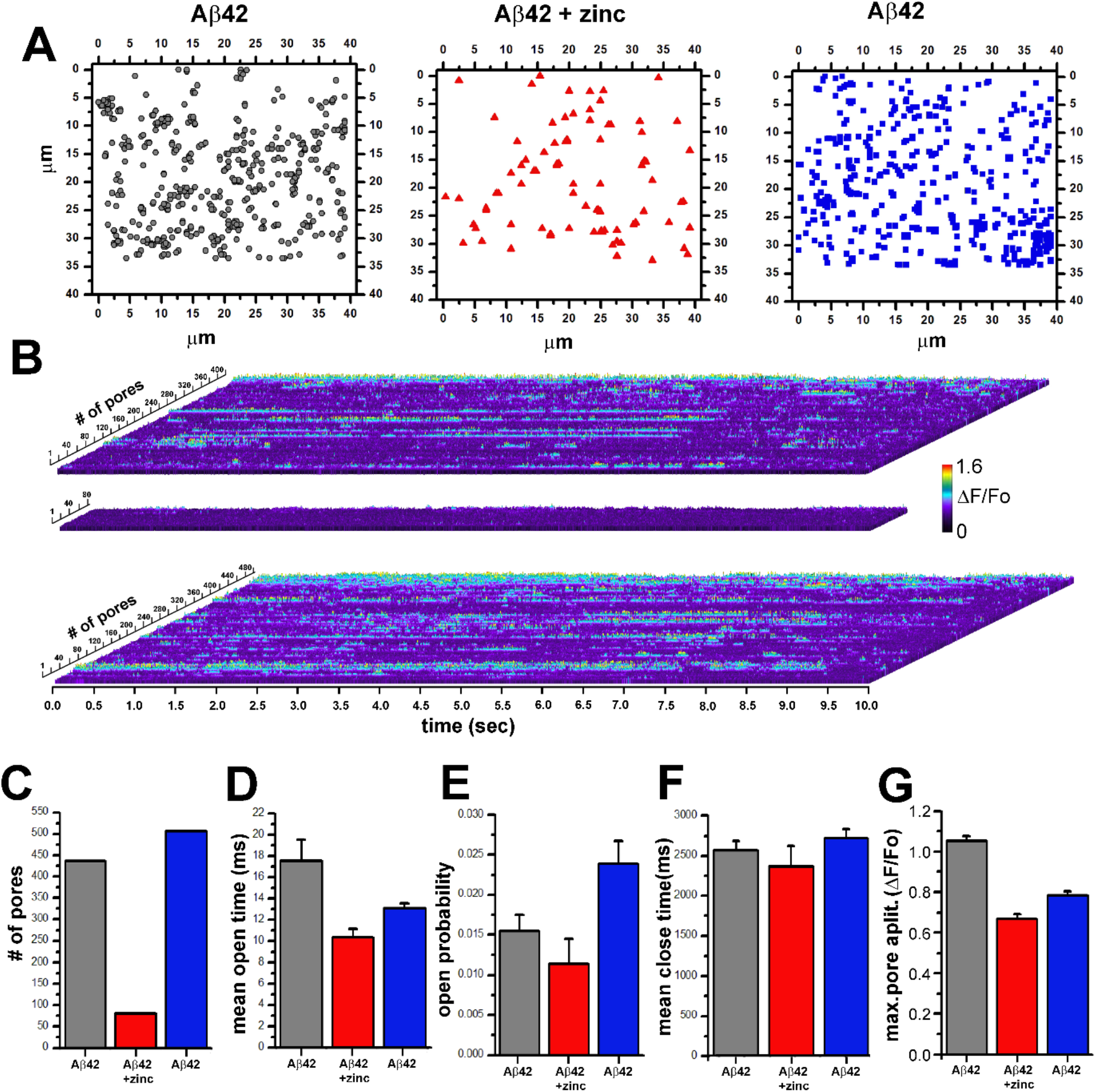
Zinc is a reversible inhibitor of Aβ42 pores. **A**) Maps of pore locations visualized in the same membrane patch in the presence of extracellular Aβ42 oligomers alone (left map), during the addition of zinc 200 μM (middle map), and after zinc removal (right map). Same concentration of Aβ42 oligomers was present during the three recordings. **B**) The corresponding color-coded images representing pores activity in the membrane patch during the entire recordings for pores shown in **A**. Top panel displays time dependent fluorescent recordings for each pore (442 pore) obtained during voltage step to −80 mV. Traces are randomly plotted from bottom to the top of the image to create a general overview of the total pore activity in the membrane patch during the 10 seconds recording. Middle panel displays Aβ42 pore (81 pores) activity recorded in the same membrane patch after addition of 200 μM zinc to the bath solution. The bottom panel depicts pore (507 pores) activity after zinc removal. **C**) The total number of pores during each recording, and the corresponding average values of mean open time **D**), open probability **E**), mean closed time **F**), and maximum amplitude **G**).

### Imaging the functional properties of Aβ40 pores

Similar to Aβ42Os, interaction of Aβ40Os with artificial and biological lipid membranes have been shown to form Ca^2+^ permeable self-gating pores with large variability in gating modality and multi steps amplitudes. However, the pores due to the two isoforms of Aβ have been rarely investigated and compared under similar experimental conditions. Here we report the results of our parallel investigation of Aβ40Os’ ability to form Ca^2+^ conducting pores. Like Aβ42Os, bath application of Aβ40Os to a voltage clamped *Xenopus* oocytes triggers the appearance of localized Ca^2+^ dependent fluorescent transients through the oocyte’s membrane patch. These Ca^2+^ transients were small and poorly detectable when the membrane potential was held at 0 mV and became clearly visible when the membrane potential was stepped to −80 mV. Experiments were performed with very similar approaches as with Aβ42Os and the Aβ40 concentration present in the bathing solution was equal (1 μg/ml). The time needed to form detectable functional pores was about 15-20 minutes, comparable to the time needed after Aβ42 oligomer addition to form pores. However, we observed that the average number of pores visualized at each recording time was smaller than those observed when Aβ42Os were added. Importantly, in contrast to Aβ42Os, we observed a general tendency of the Aβ40 pores to decrease in number during multiple consecutive stack recordings (Figure 9A). The left panel in Figure 9B shows color-coded superimposed location maps for all the pores detected during 6 consecutive stacks recorded at the same membrane patch. Panel to the right is the expanded view of the area depicted in the left panel. As for Aβ42 maps, we searched for overlapping symbols indicating the presence of persistent pores. Although we find some overlapping symbols, the overall number of persistent Aβ40 pores detected were much smaller than those present during Aβ42Os experiments, suggesting a more transient behavior of Aβ40 pores. Analysis of the pores mean open time is shown in plot Figure 9C. The corresponding average close time, averaged open probability, and averaged maximum pores amplitudes are shown in Figure 9D, 9E, and 9F respectively. Apart from the averaged close time, the values of mean open time, open probability, and maximum pores amplitudes tend to mostly decrease in each subsequent recording stacks.

**Figure 9.**
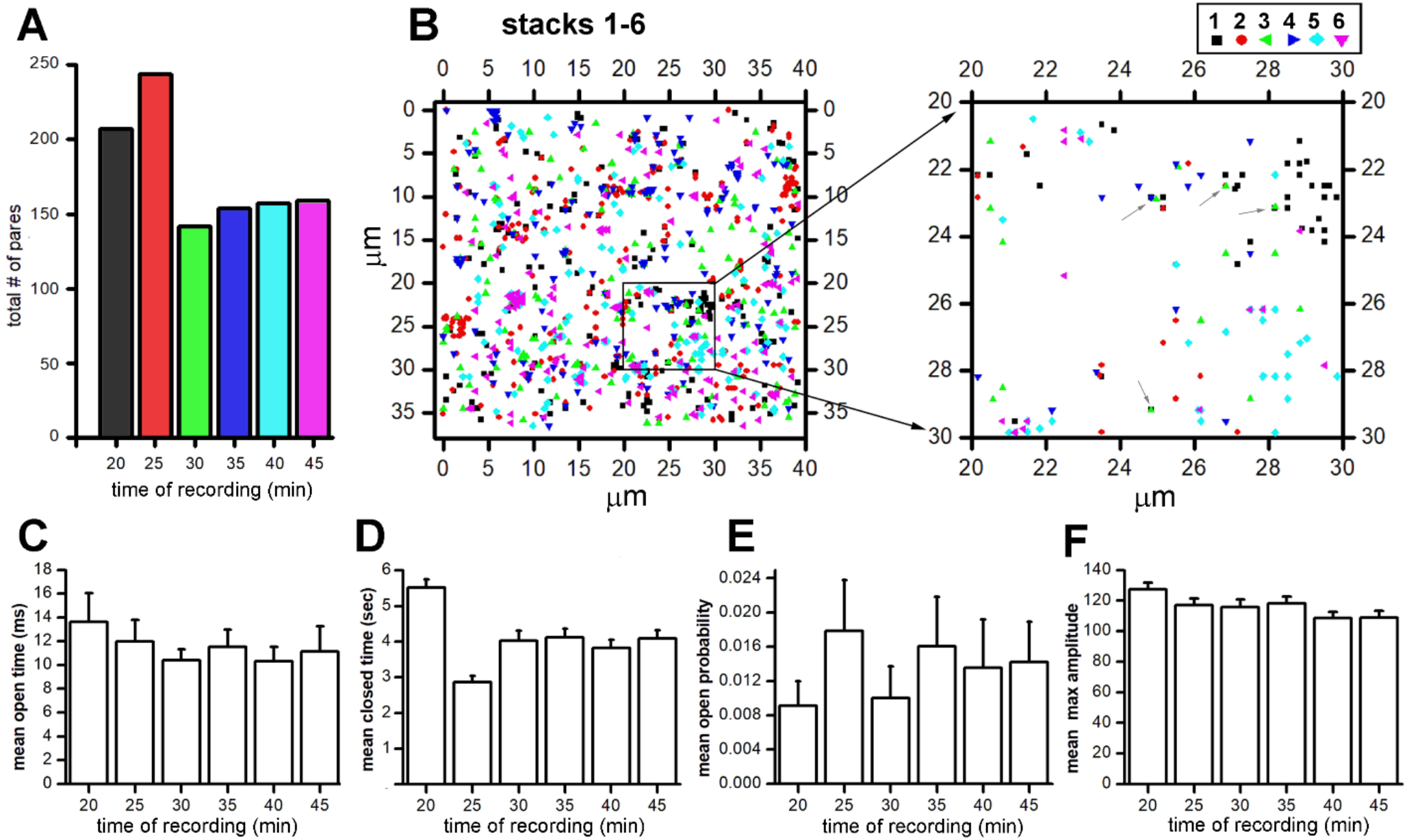
Functional Ca^2+^ permeable pores induced by bath application of Aβ40 oligomers. Bath application of Aβ40Os to *Xenopus* oocytes triggers the appearance of localized Ca^2+^ fluorescent transients through the oocyte’s membrane, similar to those seen after the application of Aβ42 oligomers. **A**) The total number of pores detected during each stack. Each recording is obtained at 5 minutes interval as indicated by the horizontal axis, and the first stack was recorded 20 minutes after Aβ40Os application. **B**) Left panel shows superimposed maps of locations of all the pores detected during 6 consecutive stacks recorded at the same membrane patch. Each color represents the pore location detected in a given stack (key shown on top of the right panel). Right panel is the expanded depiction of the area indicated in the left panel. **C**) Each bar in the plot represents the averaged mean open times for all the pores detected in each stack. Plots in **D** and **E** show the corresponding average mean close time and averaged open probability. **F** is the distribution of the mean pore maximum amplitudes in each stack.

Previous electrophysiological experiments have shown that zinc functions as a strong inhibitor of Aβ40 pores. In Figure 10, we show the analysis from three sets of experiments where the effect of zinc on Aβ40 pores’ functioning was investigated. The four traces on the top of Figure 10A are representative fluorescent traces recorded when Aβ40 was present alone, the middle set of traces are recorded after addition of 200 mM zinc, and bottom set of traces are after removal of zinc. As show in the four middle traces of Figure 10A, co-application of zinc efficiently reduces the number of detectable pores whereas zinc removal replenishes the number of detectable pores (bottom traces) as compared to control traces (top row). Plots in Figure 10B through 10E are the corresponding measurements of mean open time, open probability, mean close time, and max amplitude during this set of experiments. The cumulative reduction in pores’ number during zinc application and the recovery after zinc removal is shown by the plot in Figure10F. Overall, these data confirm the inhibitory effect of zinc on Aβ40 pores in a very similar modality as shown for Aβ42 pores. In Figure10G top three panels show pore locations in the presence of Aβ40 alone (left map), after addition of 200 mM zinc (central map), and after zinc removal (right map). Bottom row first superimposes maps of pore locations in the presence of Aβ40 before (gray symbols) and after addition of zinc (red symbols), showing mostly a lack of overlapping symbols indicating low number of common pores between the two stacks recordings (stack 1-2). In other words, the pores active in the presence of zinc are mostly newly formed pores that were not present before the zinc application. Central panel is generated by superimposing pore map from stack 1 (Aβ40 alone before zinc) with map from stack 3 (Aβ40 alone after zinc removal). The right panel shows pore locations in a 5×5 μm^2^ patch after magnification of the inset in central panel (stack1-3). The overlap of some pore locations means that while new pores have become functional after zinc removal (or during zinc application), some of the pores that were functional before zinc application are also detectable after zinc washout.

**Figure 10.**
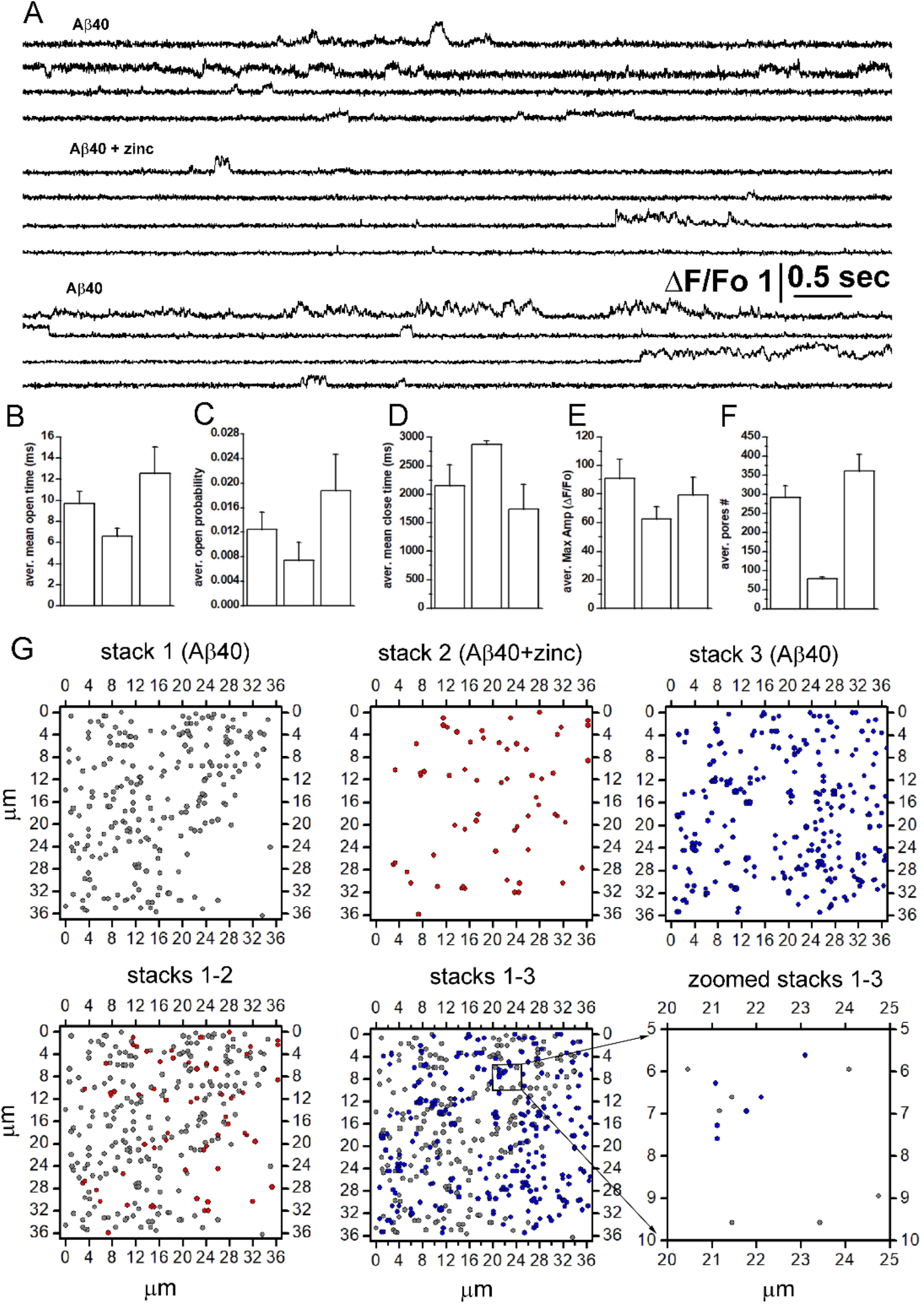
Aβ40 pores activity is reversibly inhibited by zinc. **A**) Selected fluorescent traces depicting the temporal evolution of single channels Ca^2+^ fluxes triggered by cellular exposure to Aβ40 oligomers (top set of 4 traces). Duration of pores opening are strongly reduced by co-application of zinc (200 μM middle set) whereas removal of zinc from the bathing solution restores the larger number of pores and longer open time (lower set). Plots showing global changes in functional properties of Aβ40 pores triggered by co-application zinc are shown in **B, C, D, E**, and **F. G**) The series of top three panels display location map for functional pores visualized during first recording when Aβ40 oligomers alone were present in the bathing solution (stack 1). Co-application of zinc strongly reduces the number of detectable pores in the membrane patch (stack 2) and zinc removal replenishes the number of detectable pores as shown in stack 3. The lower-left row shows superimposed maps of pore locations before and during zinc addition (stack 1-2). Central panel is generated by superimposing map from stack 1 and map from stack 3 (zinc washout) and right panel shows pore location in a 5×5 μm^2^ membrane patch after magnifying the inset in central panel (stack1-3).

## Discussion

Significant experimental evidence shows that soluble forms of Aβ aggregates lead to the upregulation of the otherwise well-controlled intracellular Ca^2+^ homeostasis, which has detrimental implications for the cell’s function and survival [13, 31-35]. Similar Ca^2+^ signaling impairments have also been observed after cell exposure to intra- or extracellular solutions containing other disease-causing oligomeric peptides, including α-Syn and PolyQ [13, 36-42]. We have previously proposed two very different mechanisms by which Aβ42Os can disrupt intracellular Ca^2+^ homeostasis; 1) by forming Ca^2+^-permeable plasma membrane pores, and 2) stimulating abnormal production of inositol 1,4,5-trisphphate (IP_3_) that triggers aberrant Ca^2+^ release from the endoplasmic reticulum (ER) through IP_3_ receptor (IP_3_R) channels. Similar mechanism of action has been proposed for the shorter isoform, the Aβ40. However, the exact molecular mechanisms underlying the impaired Ca^2+^ homeostasis, whether different isoforms use similar or different mechanisms to impair Ca^2+^ signaling, and how impairments in Ca^2+^ signaling evolve over time remain poorly understood.

Our ability to simultaneously and independently image Ca^2+^ fluxes through thousands of channels with millisecond resolutions provides a unique opportunity to investigate the formation, evolution, and function of Aβs pores in biological membranes [9, 21, 28, 43, 44]. Using our state-of-the-art imaging technique, we resolved the gating of individual Aβ42 pores in the plasma membrane of *Xenopus Leavis* oocytes and showed that several minutes after bath application of Aβ42Os, numerous localized transient Ca^2+^ influx events, representing the channel-like behavior of plasma membrane Aβ42 pores can be resolved.

In this study we further investigated the temporal evolution of Aβ42 pores formed in the plasma membrane of Xenopus oocytes over longer time, tracking each single pore over multiple recording stacks. Moreover, we performed parallel experiments imaging pores formed by Aβ40Os the 2 amino acid shorter isoform. As shown in Figure 1, automated processing of these imaging data greatly facilitated the analysis of single channel recording from the several hundreds of Aβ42 pores often recorded in each single image stack. In addition, as shown in Figure 2, we were able to track each individual pore along multiple recording stacks acquired at the same membrane patch with sub-μm spatial resolution for over a period of 90 minutes, allowing precise spatial localization of each pore over time. Analysis of data obtained from 5 different experiments with 5 different oocytes clearly indicates the tendency of Aβ42 pores to became more harmful over time (Figure 3). As shown in Figure 3A-C, the number of pores in the membrane patch increases over time as new pores are formed at later time after exposing the cell to Aβ42 oligomers. In addition, we observed that pores recorded in later time display higher mean open time, higher open probability, and higher conductance than the pores detected earlier in the experiments. The ability of *optical patch clamp* to provide high spatial resolution images of pore locations over time allows to generate pores’ location map with super resolution capability (Figure 2) [9, 23, 24, 45]. Initial visual inspection of the location of individual pores in consecutive stacks revealed the tendency of some pores to be detected only in one stack and others to be detected in as many as 6 or 7 consecutive stacks. To better understand this functional behavior, we developed an automated routine capable to link pore locations and their functional parameters over multiple stacks recording. We then performed a detailed spatiotemporal analysis of each pore detected during a single experiment consisting of 7 consecutive recording stacks (Figure 3, 4). In this particular experiment as shown in the series of plots in Figure 4A, a strong increase was observed in the number of detected pores and their respective open probability over time.

In figure 4B we show a progressive overlapping of the seven maps acquired at 5 minutes intervals.. Overlapping of these maps revealed pores that are seen only one time, either in the first or any other stacks during the duration of the experiment, that we called *transient* pores. We also observed pores that were detected in multiple stacks, that we called *permanent* pores. We then used our automated routine to link individual pore’s location within a 100 nm radius over multiple stacks (Figure 2, 4). Surprisingly, the relative number of pores are very similar, with the number of transient pores detected in each stack being constantly slightly larger. At this moment, we cannot exclude weather this functional behavior is triggered by intrinsic properties of different Aβs aggregates or by local differences in membrane lipids microdomain composition that could affect Aβs pore stability [46].

However, we can speculate that transient pores can be continuously formed as long as Aβs are available in the extracellular environment. This situation can transiently occur when a diseased neuron, rich in intracellular AβOs, release its high content of AβOs to nearby neurons by a mechanism such as apoptosis [47]. This is a transient situation after which we anticipate that only persistent pores will remain active to continue to impair intracellular Ca^2+^ homeostasis.

We speculate that the continuous rise in the function of Aβ42 pores would correlate with the progression of AD over longer duration even if neurons are transiently exposed to Aβ oligomers. Thus, the initial stage of the disease may involve the formation of small-conductance, low open probability pores that although might not immediately trigger apoptosis, but may produce chronic cell stress. This may ultimately alter the sustainable level of cytosolic Ca^2+^, which will also be compounded by the continuous presence of even high-conductance and high open probability pores that could contribute a disproportionately large Ca^2+^ influx. This assertion aligns well with the updated Ca^2+^ hypothesis of AD and aging that resulted from the deliberations of the Alzheimer’s Association Ca^2+^ Hypothesis Workgroup in 2016 [35]. This updated hypothesis specifically suggests replacing the notion of a binary state (e.g. healthy versus diseased) with the idea of a continuum of functioning/performance of a neuron. Our results from tracking cells for more than an hour after exposure to Aβ40Os Aβ42Os lend strong support to this continuum hypothesis of Ca^2+^ triggered neurodegeneration where we see continuously increasing values of mean open time, open probability, and amplitude of the persistent pores (Figures 5). We also investigated the possible relationship between the time gap at which a pore is detected in two stacks with their functional properties. The result of this analysis is shown in the series of plots in Figure 5D indicating that pore measured at longer time gap between when first detected and the second measurement, tend to have higher values in mean open time, open probability, and amplitudes. More work is needed to further test the hypothesis of continuous worsening of Ca^2+^ disruptions as the main cause for the continuum of impairments in neuronal function. While tracking Ca^2+^ changes in tissues or cell cultures over weeks and months is not possible experimentally, detailed biophysical models built on the extensive data such as presented in this paper offers a viable alternative.

In addition to the progressive rise in the toxicity at a fixed clamping voltage of −80 mV, we varied the membrane potential between 0 mV and −100 mV and found a consistent rise in not only the conductance as expected due to increase of the driving force for Ca^2+^, but also in the open probability and mean open time of pores. This is crucial as the neuronal membrane potential in resting conditions sets around this value. Thus, the higher activity of Aβ42 pores would most likely play a significant part in the observed higher intracellular Ca^2+^ concentration in neurons from AD brains (247.3±10.1 nM in adult triple transgenic AD mice versus 110.8±1.5 nM in non-transgenic adult mice) in resting conditions [48]. Such persistent high levels of Ca^2+^ would leave neurons vulnerable to impaired functioning and potentially to apoptosis in longterm. In line with this assertion, Hernandez and colleagues recently demonstrated that oral application of the compound anle138b, before or after the onset of the pathology, restores hippocampal synaptic and transcriptional plasticity as well as spatial memory in mouse model of AD by blocking the activity of conducting Aβ pores [15] suggesting that Aβs pores could play a significant role in the pathology of AD.

Zinc have been identified as the traditional inhibitor for both isoforms of Aβs pores [16, 49, 50]. Our experiments on the effect of zinc on the function of individual pores showed that in addition to strongly reducing the number of detectable pores, zinc also diminishes both the mean open time and pores amplitude. Interestingly, although a strong reduction of total number of events is observed, the average number of events per pore for pores that remain functional actually increased as expected for a typical ion channel blocker. However, zinc did not stop the formation of new pores. Furthermore, the new pores formed longer after exposing the cells to Aβ42 oligomers might still be more toxic than those formed earlier. Thus, minimizing the toxicity of already existing and new pores would require the consistent presence of zinc. We also investigated the reversibility of zinc inhibition on Aβ42 pores by imaging Aβ42 pores activity before zinc addition, in the presence of zinc, and after its removal by imaging the same membrane patch (Figure 8). After zinc removal, the number of detectable Aβ42 pores raised again reaching even higher numbers compare to the first recording (Figure 8A). Thus, the reduced gating properties of the pores were completely reverted upon zinc removal.

Together with Aβ42Os, the shorter isoform Aβ40Os have been also shown to form Ca^2+^ permeable pores. However, although up to date there is general agreement on Aβ42Os ability to form Ca^2+^ permeable pores, recent controversy has risen about the pore formation ability of Aβ40Os. Thus, we performed parallel experiments and investigated Aβ40Os’ ability to form Ca^2+^ permeable pores upon interaction with Xenopus oocytes plasma membrane. Oligomerization of Aβ40 peptides were carried out following the protocol we previously used for the oligomerization of Aβ42 peptides and experiments were performed using equal concentration of Aβ40 (1μg/ml) as for Aβ42 peptides [9]. Similar to Aβ42Os, bath application of Aβ40Os triggered localized fluorescent transients within 20 minutes after application (Figure 9A and 10A). The amplitudes and temporal evolution of these transients were very similar to those generated after application of Aβ42Os. However, although the equivalent content of Aβ40Os was used, we observed a substantially smaller number of pores in each stack recording as compared to Aβ42Os. Moreover, although we observed formation of both transient and persistent Aβ40 pores (Figure 9B), in contrast to Aβ42 pores, the total number of pores detected in later stacks decreased as function of time, as did their mean open time and amplitudes (Figure 9D and 9F). On the other hand, the open probability tended to grow over time (Figure 9E). As for Aβ42 pores and confirming what reported in literature for Aβ40 pores, addition of 200 μM zinc to the bathing solution strongly reduced the total number of Aβ40 pores detected during the recording of each consecutive stack, together with reduction of their mean open time, open probability, and pores amplitudes [9, 16, 51, 52]. However, we observed a progressively increased value of mean close time (Figure 10B - F). In this particular experiment overlapping of maps revealed that only transient pores were detected during the experiments.

In summary, our observations provide new insights into the kinetics and evolution of Aβ40 and Aβ42 pores over longer time. Our results using single channel imaging confirmed the ability of both isoforms to form Ca^2+^ permeable pores and revealed two distinct populations of Aβs pores, transient pores that are active for very brief time and persistent pores that can be recorded for much longer time. For both isoforms, we estimated that the pores lifetimes can range from < 100 ms to > 100000 ms, based on the observations of few thousands individual pores for up to 90 minutes. Although this functional behavior is share by the two Aβs isoform, the Ca^2+^ toxicity due to Aβ42Os pores tends to progressively grow overtime due to increase in pore number, amplitude, mean open time, and open probability. On the other hand, pores formed by Aβ40 display opposite feature with a time dependent rundown in Ca^2+^ toxicity. Moreover, our imaging approach would be equally useful to investigate the functional properties of other amyloid-related diseases, including Huntington’s, Parkinson’s, and prion disease, where increased membrane Ca^2+^ permeability has been observed.

## Author Contributions

S I Shah: Performed research, contributed analytical tools, analyzed data, and contributed to write the paper.

I Parker: Designed research and wrote the paper.

G Ullah: Designed research, contributed analytical tools, analyzed data, and contributed to write the paper.

A Demuro: Designed research, performed experiments, contributed analytical tools, analyzed data, and wrote the paper.

## Acknowledgments

This works was supported by NIH through grants R01 AG053988 (to AD and GU) and R01 GM065830 (to IP).

## Notes

### Competing Interest Statement

The authors have declared no competing interest.

## References

1. Hardy, J.A. and G.A. Higgins, Alzheimer’s disease: the amyloid cascade hypothesis. Science, 1992. 256(5054): p. 184.

2. Haass, C., et al., Amyloid Beta-Peptide Is Produced by Cultured-Cells during Normal Metabolism. Nature, 1992. 359(6393): p. 322–325.

3. Small, D.H., D.W. Klaver, and L. Foa, Presenilins and the gamma-secretase: still a complex problem. Molecular Brain, 2010. 3.

4. Cras, P., et al., Senile plaque neurites in Alzheimer disease accumulate amyloid precursor protein. Proceedings of the National Academy of Sciences, 1991. 88(17): p. 7552–7556.

5. Arispe, N., E. Rojas, and H.B. Pollard, Alzheimer disease amyloid beta protein forms calcium channels in bilayer membranes: blockade by tromethamine and aluminum. Proceedings of the National Academy of Sciences, 1993. 90(2): p. 567–571.

6. Lal, R., H. Lin, and A.P. Quist, Amyloid beta ion channel: 3D structure and relevance to amyloid channel paradigm. Biochim Biophys Acta, 2007. 1768(8): p. 1966–75.

7. Lin, H., R. Bhatia, and R. Lal, Amyloid beta protein forms ion channels: implications for Alzheimer’s disease pathophysiology. Faseb Journal, 2001. 15(13): p. 2433–2444.

8. Lin, H., R. Bhatia, and R. Lal, Amyloid beta-protein forms ion channels: A direct mechanism for Alzheimer’s disease pathophysiology. Biophysical Journal, 2002. 82(1): p. 17a–17a.

9. Demuro, A., M. Smith, and I. Parker, Single-channel Ca(2+) imaging implicates Abeta1-42 amyloid pores in Alzheimer’s disease pathology. J Cell Biol, 2011. 195(3): p. 515–24.

10. Ullah, G., et al., Analyzing and Modeling the Kinetics of Amyloid Beta Pores Associated with Alzheimer’s Disease Pathology. Plos One, 2015. 10(9).

11. Shah, S.I., et al., TraceSpecks: a software for automated idealization of noisy patch-clamp and imaging data. Biophysical journal, 2018. 115(1): p. 9–21.

12. Shirwany, N.A., et al., The amyloid beta ion channel hypothesis of Alzheimer’s disease. Neuropsychiatric disease and treatment, 2007. 3(5): p. 597.

13. Demuro, A., et al., Calcium dysregulation and membrane disruption as a ubiquitous neurotoxic mechanism of soluble amyloid oligomers. Journal of Biological Chemistry, 2005. 280(17): p. 17294–17300.

14. Diaz, J.C., et al., Small molecule blockers of the Alzheimer Aβ calcium channel potently protect neurons from Aβ cytotoxicity. Proceedings of the National Academy of Sciences, 2009. 106(9): p. 3348–3353.

15. Hernandez, A.M., et al., The diphenylpyrazole compound anle138b blocks A beta channels and rescues disease phenotypes in a mouse model for amyloid pathology. Embo Molecular Medicine, 2018. 10(1): p. 32–47.

16. Arispe, N., H.B. Pollard, and E. Rojas, Zn2+ interaction with Alzheimer amyloid beta protein calcium channels. Proc Natl Acad Sci U S A, 1996. 93(4): p. 1710–5.

17. Rhee, S.K., A.P. Quist, and R. Lal, Amyloid beta protein-(1-42) forms calcium-permeable, Zn2+-sensitive channel. Journal of Biological Chemistry, 1998. 273(22): p. 13379–13382.

18. Kawahara, M., et al., Alzheimer’s disease amyloid beta-protein forms Zn2+-sensitive, cation-selective channels across excised membrane patches from hypothalamic neurons. Biophysical Journal, 1997. 73(1): p. 67–75.

19. Arispe, N., J.C. Diaz, and O. Simakova, Aβ ion channels. Prospects for treating Alzheimer’s disease with Aβ channel blockers. Biochimica et Biophysica Acta (BBA)-Biomembranes, 2007. 1768(8): p. 1952–1965.

20. Kayed, R., et al., Conformation dependent monoclonal antibodies distinguish different replicating strains or conformers of prefibrillar A beta oligomers. Molecular Neurodegeneration, 2010. 5.

21. Demuro, A. and I. Parker, “Optical patch-clamping”: Single-channel recording by imaging Ca2+ flux through individual muscle acetylcholine receptor channels. Journal of General Physiology, 2005. 126(3): p. 179–192.

22. Demuro, A. and I. Parker, Imaging single-channel calcium microdomains by total internal reflection microscopy. Biol Res, 2004. 37(4): p. 675–9.

23. Demuro, A. and I. Parker, Imaging the activity and localization of single voltage-gated Ca(2+) channels by total internal reflection fluorescence microscopy. Biophys J, 2004. 86(5): p. 3250–9.

24. Demuro, A. and I. Parker, “Optical patch-clamping”: single-channel recording by imaging Ca2+ flux through individual muscle acetylcholine receptor channels. J Gen Physiol, 2005. 126(3): p. 179–92.

25. Demuro, A. and I. Parker, Optical single-channel recording: imaging Ca2+ flux through individual ion channels with high temporal and spatial resolution. J Biomed Opt, 2005. 10(1): p. 11002.

26. Demuro, A. and I. Parker, Imaging single-channel calcium microdomains. Cell Calcium, 2006. 40(5-6): p. 413–22.

27. Shah, S.I., et al., CellSpecks: A Software for Automated Detection and Analysis for Calcium Channels in Live Cells. Biophysical Journal, 2018. 114(3): p. 291a–291a.

28. Demuro, A. and I. Parker, Optical single-channel recording: imaging Ca2+ flux through individual ion channels with high temporal and spatial resolution. Journal of Biomedical Optics, 2005. 10(1).

29. Syed Islamuddin Shah, M.S., Divya Swaminathan, Ian Parker, Ghanim Ullah, Angelo Demuro, CellSpecks: A Software for Automated Detection and Analysis of Calcium Channels in Live Cells. BioRxiv, 2018.

30. Reiss, A.B., et al., Amyloid toxicity in Alzheimer’s disease. Rev Neurosci, 2018. 29(6): p. 613–627.

31. Bezprozvanny, I. and M.P. Mattson, Neuronal calcium mishandling and the pathogenesis of Alzheimer’s disease. Trends in neurosciences, 2008. 31(9): p. 454–463.

32. Demuro, A. and I. Parker, Cytotoxicity of Intracellular A beta(42) Amyloid Oligomers Involves Ca2+ Release from the Endoplasmic Reticulum by Stimulated Production of Inositol Trisphosphate. Journal of Neuroscience, 2013. 33(9): p. 3824–3833.

33. Demuro, A., I. Parker, and G.E. Stutzmann, Calcium Signaling and Amyloid Toxicity in Alzheimer Disease. Journal of Biological Chemistry, 2010. 285(17): p. 12463–12468.

34. Berridge, M.J., Calcium hypothesis of Alzheimer’s disease. Pflügers Archiv-European Journal of Physiology, 2010. 459(3): p. 441–449.

35. Alzheimer’s, A.C.H.W., Calcium Hypothesis of Alzheimer’s disease and brain aging: A framework for integrating new evidence into a comprehensive theory of pathogenesis. Alzheimer’s & dementia: the journal of the Alzheimer’s Association, 2017. 13(2): p. 178.

36. Demuro, A. and I. Parker, Cytotoxicity of intracellular aβ42 amyloid oligomers involves Ca2+ release from the endoplasmic reticulum by stimulated production of inositol trisphosphate. The Journal of Neuroscience, 2013. 33(9): p. 3824–3833.

37. Demuro, A., M. Smith, and I. Parker, Single-channel Ca2+ imaging implicates Aβ1–42 amyloid pores in Alzheimer’s disease pathology. The Journal of cell biology, 2011. 195(3): p. 515–524.

38. Magi, S., et al., Intracellular calcium dysregulation: implications for Alzheimer’s disease. BioMed research international, 2016. 2016.

39. Arbel-Ornath, M., et al., Soluble oligomeric amyloid-β induces calcium dyshomeostasis that precedes synapse loss in the living mouse brain. Molecular neurodegeneration, 2017. 12(1): p. 1–14.

40. Angelova, P.R., et al., Calcium is a key factor in α-synuclein induced neurotoxicity. J Cell Sci, 2016: p. jcs. 180737.

41. Mironov, S.L., α-Synuclein forms non-selective cation channels and stimulates ATP-sensitive potassium channels in hippocampal neurons. The Journal of physiology, 2015. 593(1): p. 145– 159.

42. Hirakura, Y., et al., Polyglutamine-induced ion channels: a possible mechanism for the neurotoxicity of Huntington and other CAG repeat diseases. Journal of neuroscience research, 2000. 60(4): p. 490–494.

43. Demuro, A. and I. Parker, Imaging the activity and localization of single voltage-gated Ca2+ channels by total internal reflection fluorescence microscopy. Biophysical Journal, 2004. 86(5): p. 3250–3259.

44. Demuro, A. and I. Parker, Imaging single-channel calcium microdomains by total internal reflection microscopy. Biological Research, 2004. 37(4): p. 675–679.

45. Wiltgen, S.M., I.F. Smith, and I. Parker, Superresolution localization of single functional IP3R channels utilizing Ca2+ flux as a readout. Biophys J, 2010. 99(2): p. 437–46.

46. Di Scala, C., et al., Common molecular mechanism of amyloid pore formation by Alzheimer’s beta-amyloid peptide and alpha-synuclein. Sci Rep, 2016. 6: p. 28781.

47. Pensalfini, A., et al., Intracellular amyloid and the neuronal origin of Alzheimer neuritic plaques. Neurobiol Dis, 2014. 71: p. 53–61.

48. Lopez, J.R., et al., Increased intraneuronal resting [Ca2+] in adult Alzheimer’s disease mice. Journal of neurochemistry, 2008. 105(1): p. 262–271.

49. Prangkio, P., et al., Multivariate analyses of amyloid-beta oligomer populations indicate a connection between pore formation and cytotoxicity. PLoS One, 2012. 7(10): p. e47261.

50. Lin, H., R. Bhatia, and R. Lal, Amyloid beta protein forms ion channels: implications for Alzheimer’s disease pathophysiology. FASEB J, 2001. 15(13): p. 2433–44.

51. Lee, J., et al., Amyloid beta Ion Channels in a Membrane Comprising Brain Total Lipid Extracts. ACS Chem Neurosci, 2017. 8(6): p. 1348–1357.

52. Lin, H., Y.J. Zhu, and R. Lal, Amyloid beta protein (1-40) forms calcium-permeable, Zn2+-sensitive channel in reconstituted lipid vesicles. Biochemistry, 1999. 38(34): p. 11189–96.

